# Birth and Death in Terminal Complement Pathway

**DOI:** 10.1101/2022.04.21.489004

**Authors:** Ashutosh Sharma, Saumya Gupta, Ajinkya Bharatraj Patil, Nagarjun Vijay

## Abstract

The cytolytic activity of the membrane attack complex (MAC) has a crucial role in the complement-mediated elimination of pathogens. Terminal complement pathway (TCP) genes encode the proteins that form the MAC. Although the TCP genes are well conserved within most vertebrate species, the early evolution of the TCP genes is poorly understood. Based on the comparative genomic analysis of the early evolutionary history of the TCP homologs, we evaluated four possible scenarios that could have given rise to the vertebrate TCP. Currently available genomic data support a scheme of complex sequential protein domain gains that may be responsible for the birth of the vertebrate *C6* gene. The subsequent duplication and divergence of this vertebrate *C6* gene formed the *C7, C8****α***, *C8****β***, and *C9* genes. Compared to the widespread conservation of TCP components within vertebrates, we discovered that *C9* has disintegrated in the genomes of galliform birds. Publicly available genome and transcriptome sequencing datasets of chicken from Illumina short read, PacBio long read, and Optical mapping technologies support the validity of the genome assembly at the *C9* locus. In this study, we have generated a >120X coverage whole-genome Chromium 10x linked-read sequencing dataset for the chicken and used it to verify the loss of the *C9* gene in the chicken. We find multiple CR1 (chicken repeat 1) element insertions within and near the remnant exons of *C9* in several galliform bird genomes. The reconstructed chronology of events shows that the CR1 insertions occurred after *C9* gene loss in an early galliform ancestor. Our study of *C6* gene birth in an early vertebrate ancestor and *C9* gene death in galliform birds provides insights into the evolution of the TCP.

## Introduction

The late or terminal complement pathway (TCP) consists of *C5, C6, C7, C8****α***, *C8****β***, *C8****γ***, and *C9* (Ricklin et al., 2010). Component *C5* is evolutionarily closely related to the *C3* and *C4* genes belonging to the α2-macroglobulin (α2M) family (Nonaka and Kimura, 2006). The MACPF domain is present in five (*C6, C7, C8****α***, *C8****β***, and *C9*) of the TCP proteins and *C8****γ*** is the only lipocalin in the TCP (Schreck et al., 2000). The MACPF domain-containing vertebrate TCP is attributed to a series of gene duplication events starting from an ancestral *C6* (Azumi et al., 2003; Hobart, 1998; Hobart et al., 1993; Mondragón-Palomino et al., 1999; Nonaka and Kimura, 2006). The ancestral *C6* and perforin, another pore-forming protein, result from an ancient duplication of the *MPEG-1* gene (D’angelo et al., 2012). While the evolution of the complement pathway has been the focus of intensive study, the emergence of TCP genes before the appearance of the vertebrate lineage is understudied (Courtney Smith et al., 2018; Nakao and Somamoto, 2016; Nonaka and Kimura, 2006; Nonaka and Yoshizaki, 2004; Sutoh and Kasahara, 2021; Zhu et al., 2005). Multiple copies of N-terminal *C6*-like genes occur in cephalochordates and urochordates (Azumi et al., 2003; Huang et al., 2008; Suzuki et al., 2002). However, these C6-like genes lack the short consensus repeats (SCR or SUSHI) and Factor I-Membrane Attack Complex (FIMAC) domain. C-terminal *C6*-like proteins (SpCRL: <u>S</u>. *<u>p</u>urpuratus* <u>c</u>omplement related protein, long-form, and SpCRS: <u>S</u>. *<u>p</u>urpuratus* <u>c</u>omplement related protein, <u>s</u>hort form) with a domain composition similar to the vertebrate TCP components occur in the purple sea urchin (*Strongylocentrotus purpuratus*). These C-terminal *C6*-like proteins contain multiple short consensus repeats (SCR or SUSHI) and Factor I-Membrane Attack Complex (FIMAC) (Multerer and Smith, 2004). However, both SpCRL and SpCRS lack the MACPF domain.

Hobart and colleagues described and compared the structures of the human TCP genes almost three decades ago (Hobart et al., 1995, 1993) and subsequently proposed a model for the evolution of the TCP components (Hobart, 1998). Large amounts of new genomic data accumulated from then allowed us to envisage four new scenarios (see **Fig. 1**) that could have given rise to the vertebrate TCP. In the first scenario, an ancestral pre-vertebrate gene containing all the six domains (TSP1, LDLa, MACPF, EGF, CCP, FIMAC) present in vertebrate *C6* underwent gene duplication followed by secondary domain loss, giving rise to the vertebrate TCP genes (see **Fig. 1A**). The second scenario involves the fusion of the N-terminal *C6*-like gene found in Cephalochordata (*Branchiostoma belcheri* and *Branchiostoma floridae*) and the C-terminal *C6*-like gene found in Echinodermata (*Strongylocentrotus purpuratus*). This gene fusion probably occurred in a hypothetical pre-vertebrate or early vertebrate ancestor which possessed both genes (see **Fig. 1B**). In the third scenario, the multiple *C6*-like genes found in pre-vertebrate species such as Urochordata (*Ciona intestinalis*) have independently gained/lost specific domains to form the various components of the TCP (see **Fig. 1C**). A fourth scenario involving sequential domain gain events resulting in a *C6* gene is also possible (see **Fig. 1D**). In scenarios 1, 2, and 4, the *C6* gene would have to undergo multiple gene duplication events to form the components of the TCP (Mondragón-Palomino et al., 1999). It is currently unknown whether a primitive pre-vertebrate TCP exists. The ancestral function of the complement system is the opsonization of pathogens, and the TCP may have evolved as a cytolytic enforcer (Courtney Smith et al., 2018).

**Figure 1:**
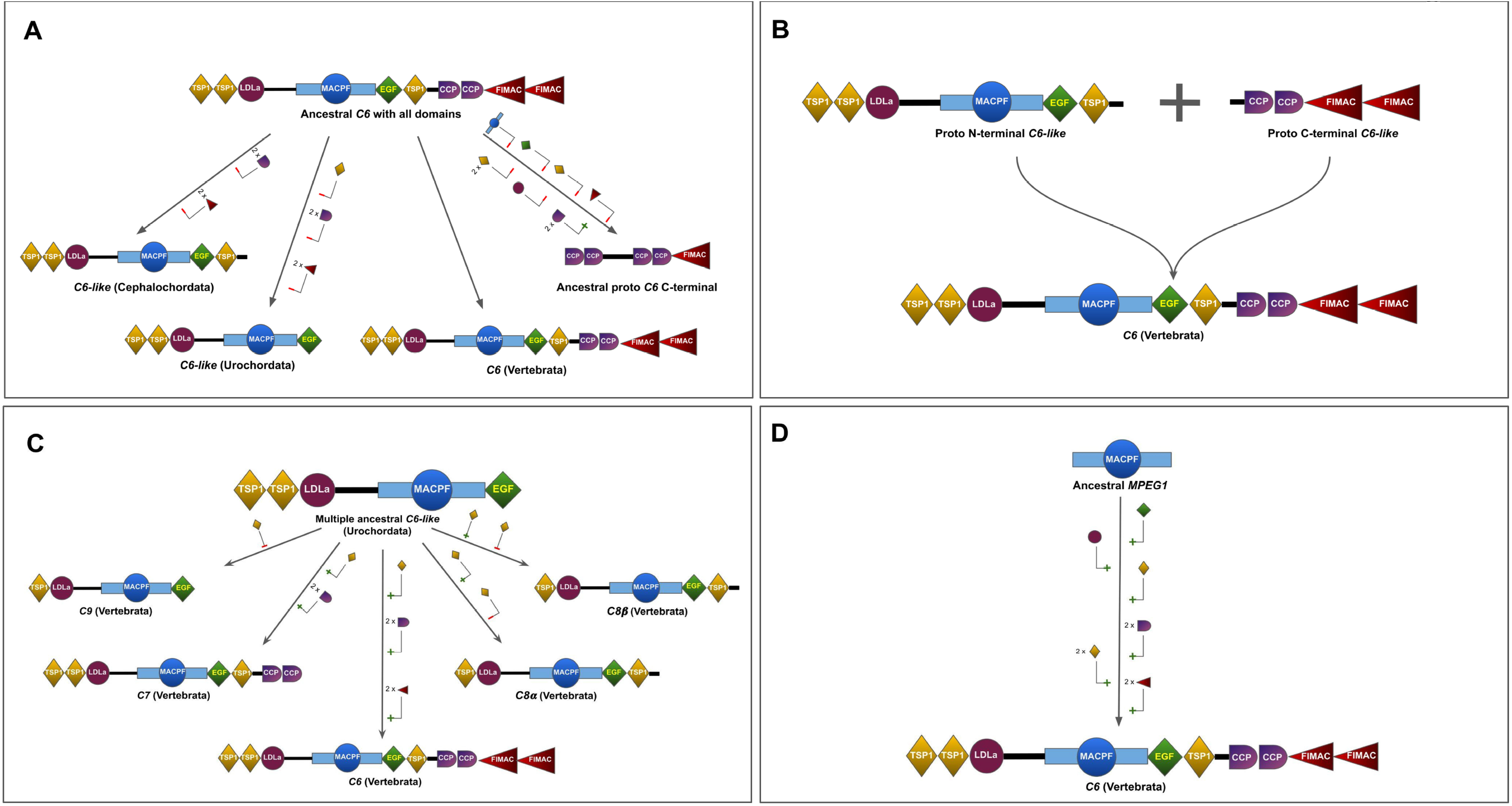
Four possible scenarios for the birth of the terminal complement pathway (TCP). **A**. An ancestral *C6* with all the domains was present in a pre-vertebrate species. The loss of 2 CCP (Sushi) domains and 2 FIMAC domains gave rise to the extant *C6-like* gene present in Cephalochordata (*Branchiostoma belcheri*), and the loss of 1 TSP1 domain, 2 CCP domains, and 2 FIMAC domains gave rise to the *C6-like* gene found in Urochordata (*Ciona intestinalis)*. Loss of 3 TSP1 domains, 1 LDLRA domain, 1 EGF-like domain, 1 FIMAC, and gain of 2 CCP domains gave rise to ancestral proto-*C6* C-terminal-like protein in Echinodermata (*Strongylocentrotus purpuratus*). **B**. The *C6* gene in vertebrates arose from the fusion of 2 genes whereby the N-terminal of *C6* derived from a proto-N-terminal *C6*-like gene from the Cephalochordata (*Branchiostoma belcheri)*, and the C-terminal of *C6* derived from a proto-C-terminal *C6-like* gene from Echinodermata (*Strongylocentrotus purpuratus*). **C**. Multiple copies of *C6*-like gene in Urochordata (*Ciona intestinalis*) independently gained different domains and gave rise to *C6, C7, C8****α***, *C8****β***, and *C9*. **D**. Ancestral *MPEG1* underwent duplication and sequentially gained 1 EGF, 1 TSP, 2 CCP and 2 FIMAC domains on its C terminal and 1 LDLa and 2 TSP domains on its N terminal to give rise to *C6* gene in vertebrates.

Among vertebrates, the genes encoding the TCP components are missing in Agnathans (jawless fishes) in contrast to Gnathostomata (jawed vertebrates) species which have a stable TCP gene content (see **Fig. 2**). The agnathan *C3b* activates the lamprey pore-forming protein (*LPFP*) to undergo polymerization to perform the cytolysis of pathogens (Matsushita, 2018; Nonaka et al., 1984; Nonaka and Takahashi, 1992; Sutoh and Kasahara, 2021; Wu et al., 2017). Within Gnathostomata, the *C5b* initiates sequential assembly of the TCP proteins resulting in the formation of membrane attack complex (MAC), a cytolytic transmembrane pore in the targeted pathogens (Lovelace et al., 2011; Müller-Eberhard, 1986; J. Tschopp et al., 1982; Jürg Tschopp et al., 1982). All three (lectin, classical, and alternative) complement pathways activate the TCP proteins that form the MAC. The MAC, which eliminates gram-negative bacteria, enveloped viruses, and parasites, relies on the well-conserved *C9* to form pore-forming multi-protein complexes (Dudkina et al., 2016; Podack and Tschopp, 1982; Xie et al., 2020). While the pores formed by the C5b-8 complex are unstable and short-lived, they can kill nucleated cells (Tegla et al., 2011). However, the relatively more stable tubular structure of the C5b-8,9n complex is required to kill gram-negative bacteria and is the target of complement evasion strategies of specific pathogens (Doorduijn et al., 2021, 2020; Joiner, 1988; Zhao et al., 2014). The *C9*-mediated killing of bacteria in the absence of C5b-8 has also been reported (Dankert and Esser, 1987). Certain viruses also target complement component *C9* as a complement evasion strategy (Bayly-Jones et al., 2017; Kim et al., 2013; Sinha et al., 2021). In humans, deficiency of the *C9* component (C9D) is a rare genetic disorder (MIM: 120940) largely prevalent in Japan and Korea, possibly due to a founder event (Khajoee et al., 2003). C9D has been linked to higher susceptibility to frequent severe infections by the pathogenic gram-negative bacteria from the Neisseria genus (*N. gonorrhoeae* and *N. meningitidis*)(Hayama et al., 1989).

**Figure 2:**
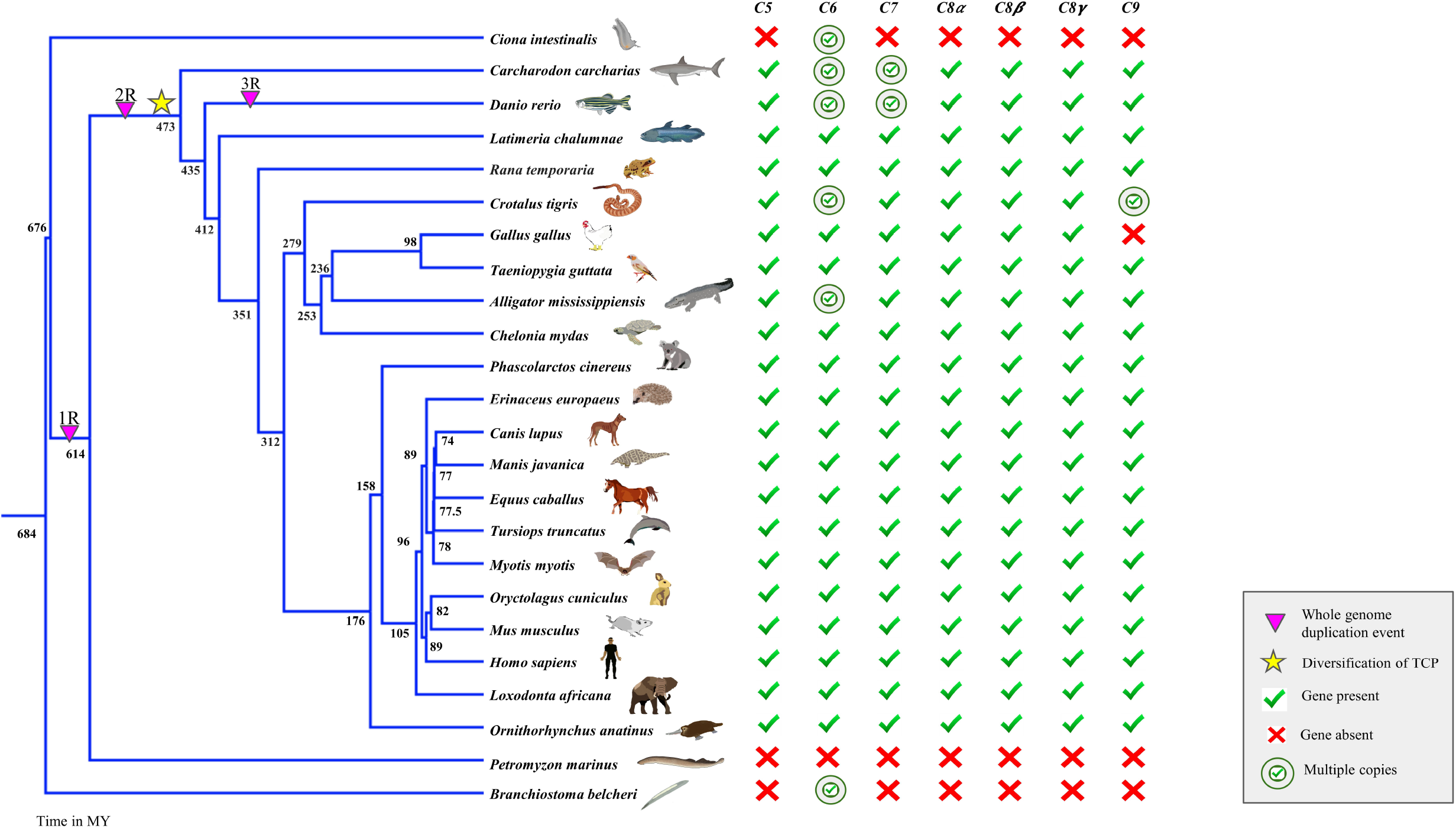
Status of terminal complement pathway (TCP) components. The phylogenetic relationship between vertebrates, Cephalochordata (*Branchiostoma belcheri*) and Urochordata (*Ciona intestinalis)* representative species. The time-calibrated phylogenetic tree is downloaded from the TimeTree website and visualized in FigTree. Whole-genome duplication (WGD) events during vertebrate evolution 1R, 2R, and 3R are shown with a filled magenta-colored triangle and indicate the first, second, and third rounds of WGD, respectively. The symbol of a star indicates the diversification of the TCP components. The right tick in green indicates the presence of genes, while the X symbol in red indicates the absence/loss of genes. The green encircled tick shows the presence of multiple copies of the same gene.

Early studies of the complement pathway components in chicken claimed to have detected the *C9* protein (Barta and Hubbert, 1978; Lynch et al., 2005). However, subsequent studies which searched genome assemblies could not find the *C9* gene in the chicken, while orthologs could be identified in mammals and amphibians (Dodds and Matsushita, 2007; Nonaka and Kimura, 2006). Due to the challenges in conclusively establishing gene loss, especially in birds (Mikrou and Zarkadis, 2010; Rohde et al., 2018), it is unknown whether the *C9* gene is intact or not in the chicken genome. The chicken genome served as a representative for the avian clade until the sequencing of more bird genomes (Zhang et al., 2014b). Hence, the inability to find *C9* in the chicken genome was viewed as a lack of *C9* in all Aves (Nonaka and Kimura, 2006). Recently, *C9* has been found in the genomes of other birds (Li et al., 2021). Despite the availability of large genomic datasets in galliform birds, it is currently unclear whether the *C9* gene is lost in chicken or simply missing from the genome assembly.

In this study, we compare the genomes of species from Cephalochordata, Urochordata, and Vertebrata to identify the set of genes homologous to components of the TCP. Based on our comparative analysis, we evaluate support for each of the possible scenarios for the emergence of the TCP. We propose the most likely sequence of events involving gene duplication, fusion, and domain gain that probably gave birth to the ancestral *C6* gene. This ancestral *C6* gene subsequently evolved to form the vertebrate TCP (*C7, C8****α***, *C8****β***, and *C9*) through gene duplication and domain loss events (Mondragón-Palomino et al., 1999; Ni and Gilbert, 2017). Although a few components of the TCP have experienced lineage-specific duplication events, no instance of gene loss is known within Gnathostomata. Surprisingly, we discovered the disintegration of the complement *C9* gene in the genomes of galliform birds. In contrast to galliform birds, the *C9* encoding gene is widely conserved across more than 338 of 362 considered vertebrate species spanning mammals, birds, non-avian reptiles, amphibians, and fishes. Thus, we illustrate that even a widely conserved gene such as terminal complement component *C9* with important immune functions can undergo lineage-specific gene loss. The comparative genomic analysis spanning >600 million years undertaken in our study helps reconstruct the process of birth and death in the TCP.

## Materials and methods

### CLANS analysis of protein domains of TCP genes

We used the CLANS software to perform sequence similarity-based clustering of all TCP protein domains except the commonly available EGF_like domain to trace the history of domain gain/loss and gene fusion in TCP components (see **Supplementary_information/CLANS_results**). We downloaded all annotated protein sequences for the thrombospondin type 1 domain (TSP_1 PF00090), low-density lipoprotein receptor domain class A (LDLRA PF00057), membrane attack complex/perforin domain (MACPF PF01823), complement control protein modules or SUSHI repeats (CCP PF00084), factor I / membrane attack complex (FIMAC IPR003884), and kazal-type serine protease inhibitor domain (Kazal_1 PF00050, Kazal_2 PF07648, Kazal_3 PF18434) from UNIPROT. We provided the downloaded protein sequences from UNIPROT as input to the CLANS java application with an e-value cut-off of 1e −4 and p-value of 0.05 to cluster the blast hits. After getting the CLANS output, we visualized it in the CLANS GUI. The clusters identified by CLANS were manually annotated based on the most common protein in that cluster.

### Collation of *C9* orthologs and annotation validation

The NCBI and Ensembl databases have annotations for 1-to-1 orthologs of the *C9* gene for ∼300 species covering all classes of the vertebrate subphylum except Agnatha (see **Supplementary Table S1, S2**). For the detailed study of *C9* evolution, we screened representative genomes of additional species using a blastn search with the *C9* gene sequence of closely related species as a query (see **Supplementary Table S1**). The open reading frame sequences of the ∼250 non-bird species were grouped at the order taxonomic level for subsequent molecular evolutionary analysis. When an order contained less than three species with sequenced genomes, we grouped it with its phylogenetically nearest order, e.g., Monotremata is grouped with Marsupials and Coelacanths with Chondrichthyes. Surprisingly, we could neither find annotations nor genomic sequences corresponding to an intact *C9* gene in species belonging to the order Galliformes in birds. Hence, to ascertain the conservation of the *C9* gene in other bird species, we screened the genomes of ∼110 bird species covering all the 42 bird orders described in the Clements Checklist of Birds of the World 2021 (see **Supplementary Table S1**).

The annotation for each copy of the *C9* gene was verified to correct partial sequences, exon number inconsistencies, sequence length variation, and incorrect open reading frame (ORF). Using the functionally validated and well-studied sequence of the *C9* gene from the human genome as a query to the blast program, we inspected the genome assemblies and short-read datasets of various species for the initial annotation validation. Conserved synteny in the vicinity of the gene is an important consideration while identifying 1-to-1 orthologs of genes from multigene families. Hence, we compared the synteny using five flanking genes upstream and downstream of the *C9* gene (see **Supplementary_information/C9_synteny_Screenshots** and **Supplementary Table S3**). The multiple sequence alignment of the open reading frame sequences at the codon level was performed using Guidance (v2.02) (Sela et al., 2015) with PRANK as the aligner. These alignments were carefully scrutinized by visual inspection and were subsequently verified using a blast search of short-read sequencing datasets in case of any discrepancies. Sequence alignments are available in **Supplementary_information/Alignment_and_Species_tree**.

### Molecular evolutionary analyses

We evaluated the multiple sequence alignments of the *C9* sequences from groups roughly corresponding to the taxonomic level of “order” for sequence saturation using the index to measure substitution saturation (Iss) described in Xia et al. (Xia et al., 2003) and implemented in the DAMBE program (v7.3.2) (Xia, 2013) (see **Supplementary_information/Sequence_saturation_test**). We downloaded a time-calibrate species tree for each group from the TimeTree website (see **Supplementary_information/Alignment_and_Species_tree**). Lineage-specific signatures of relaxation/intensification of selection were identified using branch tests in codeml from the PAML (Yang, 2007) package and the program RELAX from the HyPhy package (Kosakovsky Pond et al., 2005) (see **Supplementary_information/PAML_branch_model, RELAX** and **Supplementary Table S4, S5**, and **S6**). Positively selected sites were also identified using the site model in PAML (see **Supplementary_information/PAML_site_model** and **Supplementary Table S7**). Sites with signatures of selection in specific lineages were identified using the branch-site method available in the aBSREL (Smith et al., 2015) program (see **Supplementary_information/aBSREL**). The magnitude of gBGC in each species was also quantified using the mapNH program from testNH(v1.3.0) (Romiguier et al., 2012) and phastBias from PHAST(v1.6) (Hubisz et al., 2011) (see **Supplementary_information/gBGC**). Repeat regions were identified using the RepeatMasker webserver (Smit et al., 1996) (see **Supplementary_information/Repeatmasker_results**).

### Verification of gene disrupting changes in Galliformes

The previously published five-pass strategy to verify gene loss has conclusively established the loss of *PLGRKT* (S. Sharma et al., 2020) and *COA1*/*MITRAC15* (Sharad Shinde et al., 2021) genes in Galliformes. We used this same procedure to verify the loss of the *C9* gene in galliform birds. In brief, we searched the genome assembly and short-read genomic and transcriptomic datasets of chicken using the blast search tool (see **Supplementary_information/blastn_results**). The intact *C9* gene sequences of several species that are phylogenetically close to galliform birds were used as a query. We ascertained the presence/absence of individual exons in chicken by relying upon this initial blast search. Subsequently, we screened genome sequencing data of other galliform birds to evaluate the presence/absence of *C9* exons. We analyzed the shared loss of *C9* exons along the galliform phylogeny to validate further and identify the timeline of gene loss. We relied upon publicly available datasets of long-read PacBio sequencing and optical mapping to rule out the possibility of genome assembly artifacts and sequencing bias. Additionally, we have generated a ∼120X coverage 10x Chromium linked-read whole genomic dataset to validate the chicken genome assembly (**Supplementary Table S8**).

The PacBio long-read sequencing data available for the chicken was mapped to the GRCg6a genome of the chicken using the bwa mem (v0.7.17-r1188) read mapper (Li, 2013). The resultant alignments in the bam format were visualized in the Integrative Genome Viewer (IGV v2.5.2) browser (Thorvaldsdottir et al., 2013). A tiling path of long reads spanning the genomic region containing the remnants of the *C9* gene and flanking genes on both sides were also visualized in the UCSC genome browser (see **Supplementary_information/Fig6**). We obtained the chicken optical mapping data provided by the Vertebrate genome project from the GenomeArk (https://vgp.github.io/genomeark/Gallus_gallus/Bionano). The optical mapping data was aligned to the chicken genome using OMBlastMapper (Leung et al., 2017b) and visualized using the OMview option in OMTools (see **Supplementary_information/Chicken_optical_mapping_data_screenshot**) (Leung et al., 2017a).

### Chromium 10x linked-read data generation

We purchased chicken (broiler breed) meat from an FSSAI (Food Safety and Standards Authority of India) licensed shop in Bhopal, Madhya Pradesh, India. Whole genomic DNA was extracted from the muscle tissue of the chicken using QIAGEN’s Blood and Tissue Kit and eluted in 70 ul elution buffer. The quality of extracted DNA was evaluated using 1% Agarose gel, Nanodrop, and Qubit(tm) 3 Fluorometer. The integrity of the DNA fragments was assessed using Agilent 2200 TapeStation with High Sensitivity D1000 ScreenTape. After validating the fragment size peak above 1500 bp, the sample was used to prepare a 10x Chromium library with a Chromium controller machine as per the manufacturer’s protocol. The prepared library fragments of around read length 150 bp were sequenced using Illumina NovaSeq 6000 with an insert size of 300 bp. The 10x .fastq files obtained after the bcl2fastq command were preprocessed using proc10xG (https://github.com/ucdavis-bioinformatics/proc10xG) to get GEM-barcode whitelist and other QC parameters for the reads. The linked-read datasets were mapped to the chicken genome (GRCg6a) using the Longranger pipeline (Zheng et al., 2016). We assessed the validity of the chicken genome assembly in the vicinity of the *C9* gene by visual inspection of this alignment in the IGV browser. We selected two tags to visualize 10x linked reads in IGV. (1) the BAM barcode tag “BX,” which is a chromium barcode sequence that is error-corrected and confirmed against a list of known-good barcode sequences, and (2) the BAM phasing tag “MI,” which is a global molecule identifier for the molecule that generated this read.

### Evaluating the transcriptional status of *C9*

To assess the transcriptional status of *C9*, we analyzed public transcriptomic datasets by mapping the RNA-seq reads to the genome assemblies using the STAR (v2.7.0d) read mapper (Dobin et al., 2013) and visualizing alignments using the IGV browser (Thorvaldsdottir et al., 2013). In the case of galliform birds, we screened all available tissues to assess the transcriptional status of *C9*. However, in other species with an intact *C9*, we required that the gene be expressed in at least one tissue. For consistent representation across tissues and species, we used two different views. (1) Positions of all annotated exons of *C9* or those identified using blast search are shown as a bed record with RNA-seq read alignments. (2) Zoomed-in view of the first and last exons of *C9* is shown along with the adjacent genes on both sides. In the absence of any transcriptional activity, images of the adjacent genes are taken as a positive control.

## Results

### The emergence of the terminal complement pathway

The genes encoding all the TCP components have conserved one-to-one orthologs across most vertebrate species (see **Fig. 2** and **Supplementary_information/Fig2**). We consider four possible scenarios (see **Fig. 1**) for the emergence of the vertebrate TCP and interrogate genomic datasets to evaluate the support for each scenario. The first scenario suggests that all TCP components result from secondary protein domain loss after an ancestral gene containing all six vertebrate TCP domains (TSP1, LDLa, MACPF, EGF, CCP, and FIMAC) underwent gene duplication (**Fig. 1A**). Currently available genomic datasets from invertebrate species lack ancestral TCP homologs with a protein domain composition matching the vertebrate *C6*. The ancestral TCP homologs may have given rise to the intermediate *C6*-like genes through secondary domain loss. For instance, *C6*-like genes which resemble the N-terminal are present in Cephalochordata (*Branchiostoma belcheri*) and Urochordata (*Ciona intestinalis*). *C6*-like genes corresponding to the C-terminal half occur in Echinodermata (*Strongylocentrotus purpuratus*). To ascertain the source of these intermediate forms, we used sequence similarity-based clustering analysis in the CLANS software to identify potential donors of each protein domain. Clustering of MACPF domain-containing proteins revealed the presence of this domain in bacteria, fungi, protozoa, plants, and animals. A closer inspection of the cluster containing animal sequences shows one large cluster with perforin (*PRF1*) homologs and distinct but closely placed groups for *C6, C7, C8*, and *C9*. The *C6*-like proteins from Cephalochordata (*Branchiostoma belcheri*) and Urochordata (*Ciona intestinalis*) are between *PRF1* and *C6*-*C9* but closer to the *C6*-*C9* cluster (see **Supplementary_information/CLANS_results/MACPF_domain_containing_proteins**). A detailed study of the evolutionary history of perforin (*PRF1*) and its homologs based on a comparison of domain composition suggests that an ancient *MPEG-1* gene duplication produced the common ancestor of *PRF1* and *C6* (D’angelo et al., 2012). Our sequence clustering analysis of MACPF domain-containing proteins is consistent with a common origin for *PRF1* and *C6*-*C9* genes.

Sequence similarity-based clustering of LDLRA domain-containing proteins from Pfam shows that *C6*-*C9* is close to a cluster of uncharacterized proteins that contain the TSP1 and LDLRA domains (see **Supplementary_information/CLANS_results/LDLRA_domain_containing_proteins**). Uniprot annotation for some of the proteins in this cluster is *HMCN-1* (Hemicentin-1), and others as SCO-Spondin-like. Although detailed characterization of these proteins is limited, we find an RNA-seq supported complete open reading frame (Accession# XM_036513019.1) in East Asian common octopus (*Octopus sinensis*). Several other Mollusca species (including *Crassostrea gigas*) have homologs for this gene, with no clear vertebrate orthologs. Therefore, the intermediate *C6*-like forms containing the N-terminal half are probably the result of a gene fusion between a *MPEG-1*-like gene and this SCO-Spondin-like gene.

We failed to find *C5, C6, C7, C8****α***, *C8****β***, *C8****γ***, and *C9* gene orthologs in the genomes and short-read sequencing data of sea lamprey (*Petromyzon marinus*) and inshore hagfish (*Eptatretus burgeri*). We did not find any orthologs of these genes in three more agnathan species (*Lethenteron camtschaticum, Lethenteron reissneri*, and *Entosphenus tridentatus*) with high-quality genomes (see **Supplementary_information/blastn_results/Agnatha**). Our comparative genomic analysis and protein domain sequence similarity-based clustering failed to support the first scenario and suggest that the vertebrate multi-domain *C6* protein was most likely born in an early vertebrate ancestor.

The second scenario involves a gene fusion event between two ancestral genes with a domain composition resembling the N-terminal and C-terminal regions of the vertebrate *C6 (***Fig. 1B**). The exons length, phase, sequence content, and domain order of the Cephalochordata gene corresponding to the N-terminal match the vertebrate C6. However, the Echinodermata (*Strongylocentrotus purpuratus*) SpCRS gene corresponding to the C-terminal does not match the vertebrate *C6*’s exons length or phase but matches the domain order. FIMAC domain sequence similarity-based protein clustering found the SpCRS gene close to the *C6*-*C7* cluster (see **Supplementary_information/CLANS_result/FIMAC_domain_containing_proteins**). The SpCRS gene differs from the C-terminal of vertebrate *C6* as it contains 4 CCP and 1 FIMAC/KAZAL domain. The two C-terminal CCP domains near to FIMAC/KAZAL domain in the SpCRS gene are encoded by single exons similar to RCA (Regulators of Complement Activation) proteins. However, two exons encode the first CCP domain in vertebrate *C6*, and a single exon encodes the second CCP. Hence, the distribution of the CCP domains over exons does not match between the SpCRS gene and vertebrate *C6*.

In the CLANS results of CCP domain-containing proteins, we find the multiple CCP domain-containing *CSMD2* (CUB and Sushi multiple domain-2) genes cluster near the *C6* cluster (see **Supplementary_information/CLANS_result/CCP_domain_containing_proteins**). At the C-terminal of *CSMD2*, we see two consecutive CCP domains with the same exon phase and domain distribution as vertebrate *C6*, but this gene does not have any FIMAC/KAZAL domain. It is possible that the purple sea urchin had a hypothetical C-terminal *C6*-like gene with the same CCP domain structure as *CSMD2/*vertebrate *C6* and also contained a FIMAC/KAZAL domain. This hypothetical gene may have fused with the N-terminal *C6*-like gene of Cephalochordata. In the vertebrate *C6* gene, the FIMAC/KAZAL spans three exons, with the first FIMAC present on two exons and most of the second FIMAC on a single exon (Hobart et al., 1993). The FIMAC/KAZAL domain in purple sea urchin spans only one exon and may have duplicated just before the gene fusion event. After the duplication, the second FIMAC/KAZAL domain may be split into two different exons. Hence, we find support for the second scenario in which either the SpCRS gene or the hypothesized purple sea urchin gene may have fused with the N-terminal *C6*-like gene of Cephalochordata.

The third scenario suggests several independent domain gain/loss events after duplication of a *C6*-like gene in Urochordata. Several copies of *C6*-like genes occur in the *Ciona intestinalis*, and *Branchiostoma belcheri* genomes and provide the necessary starting point for independent domain gains (see **Fig. 1C** and **Supplementary Table S9**). However, the vertebrate TCP components are unlikely to be derived from independent domain gain/loss events as all TCP components have matching exon lengths and phases for the shared regions. The large number of domain gains required by this scenario does not provide the most parsimonious explanation. Moreover, the gene order in the vicinity of TCP components suggests (based on a “Strict” Q-score criteria in http://ohnologs.curie.fr/) that *C6, C7*, and *C9* are ohnologs of *C8****α*** and *C8****β*** resulting from a whole-genome duplication. The genes *C8****α*** and *C8****β*** result from segmental duplication and occur beside each other, flanked by *FYB1* and *DAB2* genes. Similarly, the *C9* gene is flanked by *FYB2* and *DAB1* genes. The genes *C6* and *C7* are also located near *PRKAA1*, while C8a, C8p are near the *PRKAA2* gene. Previous studies also support TCP components arising from duplication events (Mondragón-Palomino et al., 1999).

The fourth scenario posits that this initial MACPF domain-containing gene has sequentially gained EGF, TSP1, LDLa, CCP, and FIMAC domains to form the vertebrate *C6* gene (see **Fig. 1D**). However, the source of these acquired domains is unknown. Sequence similarity-based clustering supports a fusion of SCO-Spondin-like gene at the N-terminal of MPEG-1-intermediate gene. However, the EGF and the TSP1 at the C-terminal result from sequential domain gain events. Subsequent fusion of the ancestral proto-*C6* originating from Echinodermata (*Strongylocentrotus purpuratus*) at the C-terminal of the C6-like gene leads to the vertebrate *C6*. Hence, we find support for scenario-2 (at least two gene fusion events) and parts of scenario-4 (at least two sequential domain gain events). Based on the well-supported events, we propose a new scheme for the emergence of vertebrate *C6*, which subsequently formed the vertebrate TCP. While the components of the TCP are conserved among vertebrates, the *C9* gene appears to be missing in the chicken. Hence, we investigated the detailed evolutionary history of the *C9* gene by analyzing the gene order at the genomic locus containing *C9* in several representative vertebrate species and validated the genome assembly.

### Conservation of *C9* syntenic region and exon structure variation

The genomic regions flanking the *C9* gene have a broadly conserved synteny (see **Fig. 4, Supplementary_information/Fig4** and **Supplementary Table S3**), with the left flank containing *DAB2, PTGER4*, and *TTC33* and the right side containing *FYB1, RICTOR*, and *OSMR*. While most of these genes are nearby (<50Kb), the region between the genes *DAB2* and *PTGER4* is ∼1Mb in the human genome and has a few annotated lncRNA (*LINC02104, LINC00603*, and *LINC00604*) and ncRNA (*ENSG00000287597, ENSG00000283286, ENSG00000285616*, and *ENSG00000285552*). In the human genome, the *C9*-containing region is part of the cytogenetic band 5p13.1, which includes the genes *C7* and *C6*. Although the Coelacanth (*Latimeria chalumnae*) genome is fragmented, most genes (except for *TTC33* and *PTGER4* located on a different scaffold) from this synteny block are assembled in the correct order in a single scaffold. The *OSMR* gene results from a duplication event originating from the *LIFR* gene located adjacent to *OSMR* but in the opposite orientation (Malaval et al., 2005). The *LIFR* gene duplication has not occurred in the great white shark lineage. However, this synteny block from the *RICTOR* gene located beside *LIFR* (an *OSMR* paralog) until *TTC33*, including the ∼1Mb gene desert, is conserved in the great white shark (*Carcharodon carcharias*). In contrast to the conserved synteny seen in most vertebrate species, we noticed that the gene order in the zebrafish (*Danio rerio*) was different (see **Fig. 4** and **Supplementary Table S3**). Although the gene *DAB2* is located upstream from *C9*, the subsequent gene order has changed to *HSPB15* and *MYHZ1*. The genes downstream from *C9* in the zebrafish are *GAS1B, FBXW2*, and *NCS1A*. Other species that share the third round of whole-genome duplication (3R-WGD) with the zebrafish also have the *DAB2* gene beside *C9*. However, the other genes in the vicinity of *C9* are heterogeneous in various 3R-WGD fish species (see **Supplementary_information/C9_synteny_Screenshots** and **Supplementary Table S3**). Notably, the non-teleost reed fish (*Erpetoichthys calabaricus*), which does not share the 3R-WGD, has the same gene order as Coelacanth and other tetrapods (see **Supplementary Table S3**). Since teleost fish have undergone the 3R-WGD, they possess additional copies of many complement pathway genes (Nakao et al., 2011). However, the overall organization of the complement pathway of teleosts is comparable to other bony fishes (Najafpour et al., 2020; Nakao et al., 2006).

Before the emergence of mammalian species, the *C9* gene had 11 coding exons in fishes (Li et al., 2007), amphibians, turtles, lizards, snakes, and birds (see **Fig. 4**). Both copies of the *C9* gene result from a tandem duplication in the order Squamata (mainly snakes) and have 11 exons each. The remnants of the *C9* gene following a gene loss event in the order of Galliformes also have 11 exons. Even in marsupial species such as the platypus (*Ornithorhynchus anatinus*), 11 coding exons are present. However, several species have only ten coding exons within mammals due to a stop codon in the tenth exon. Interestingly, an 11^th^ coding exon is regained in carnivores, bats, camels, and most primates due to independent substitutions.

### Duplication and degeneration of *C9* in Squamata

Two intact copies of the *C9* gene could be identified adjacent to each other in the genomes of several species of the order Squamata (see **Fig. 4**). While the intact duplicate copies are found mainly in snakes, the anole lizard (*Anolis carolinensis*) also has two intact copies of *C9*. Although the two copies are distinct and appear in several closely related species, the level of sequence divergence between them is less than ∼15%. Phylogenetic analysis of both copies of *C9* from squamate species, the single intact copy of *C9* from outgroup species, *C8****α*** and *C8****β***, revealed (see **Fig. 5** and **Supplementary_information/Fig5**) separate clustering of *C8****α*** and *C8****β*** but no clear separation of the two copies of *C9*. Bootstrap support for the nodes separating *C8****α*** from *C8****β*** and *C9* copies from *C8* copies are strong (see **Supplementary_information/Fig5**). However, the bootstrap support for the internal nodes within the *C9* cluster is weak.

Nonetheless, both copies of *C9* appear to be robustly expressed in the anole lizard (see **Supplementary_information/IGV_screenshots/representative_species_RNAseq**). While the copy of the *C9* gene closer to the *DAB2* gene, referred to as *C9A*, is intact in the Indian python (*Python molurus*) genome, the other copy closer to *FYB1*, referred to as C9B, has degenerated. Similarly, the copy closer to *DAB2* is intact in several species of lizards (*Sceloporus undulatus, Pogona vitticeps, Lacerta agilis, Zootoca vivipara, Podarcis muralis*, and *Gekko japonicus*), and the copy closer to *FYB1* has degenerated (see **Supplementary_information/Fig5**). Remains of a second copy of the *C9* gene besides an intact copy are also present in the genomes of several species of birds, insectivores, and Xenarthra (see **Supplementary Table S10**).

### Verification of *C9* gene loss by validating genome assembly correctness

Conclusively establishing the loss of a gene is confounded by errors in the genome assembly and the prevalence of biases in sequencing technologies. The chicken genome assembly is relatively high quality and provides an accurate picture of the actual sequence. Short-read datasets also support the patterns of exon loss seen at the *C9* locus in the chicken genome assembly (see **Supplementary_information/blastn_results**). However, local errors in the genome assembly due to collapsed repeats, incorrect scaffolding, and gaps are a genuine possibility. Hence, we rely upon multiple genomic technologies to verify the correctness of the genome assembly. First, we confirmed the correctness of the chicken genome assembly using PacBio long-read sequencing data based on a series of overlapping >10kb reads that span the entire region of the *C9* gene remnants up to adjoining genes (*FYB1* and *DAB2*; see **Fig. 6, Supplementary_information/Fig6** and **Supplementary_information/Chicken_PacBio_screenshot**). Next, we used publicly available optical mapping data to verify the long-range contiguity of the genomic locus containing the remnants of the *C9* gene (see **Supplementary_information/Chicken_optical_mapping_data_screenshot**). Finally, we rely upon chicken whole-genome Chromium 10x linked-read data generated as a part of the current study to verify the correctness of the genomic region between *FYB1* and *DAB2* (see **Supplementary_information/Chicken_10X_linked_read_screenshot**). We found that all three genomic technologies with long-range contiguity information support the validity of the genome assembly at the *C9* locus. Hence, we can rule out the possibility of a genome assembly error at the *C9* locus.

### The gradual disintegration of complement *C9* gene in galliform birds

Genomic and transcriptomic assays consisting of high-quality genomes, Illumina short-read, PacBio long-read, Chromium 10x linked-read, optical mapping, and RNA-seq data from more than twenty tissues support the loss of the *C9* gene in the chicken genome. To track the evolutionary chronology of events involved in the loss of the *C9* gene, we screened the genomes of 26 species of Galloanserae birds. Interestingly, all 18 screened galliform bird genomes have lost the *C9* gene compared to an intact *C9* in all eight screened anseriform genomes (**Fig. 7, Supplementary_information/blastn_results** and **Supplementary_information/IGV_RNAseq_screenshots/Galliformes**). Hence, *C9* gene loss appears to have occurred after the split of galliform and anseriform lineages sometime between 68 and 80 Mya in an early galliform ancestor. Careful examination of the individual exons of the *C9* gene genomes and short-read datasets revealed that exon-8 is entirely missing in all 18 galliform species. The remaining nine exons show differing levels of sequence degeneration. Exons one, seven, nine, and ten are intact in only one of the eighteen species considered. Exons four and six are intact in 16 species and completely missing in 2 other species. Exons five and eleven are intact in four and five species, respectively. Exon-2 is entirely missing in *Numida meleagris* and disrupted by a chicken repeat 1 (CR 1) insertion in *Coturnix japonica*. This CR1 insertion in exon-2 is unique to the Coturnix lineage and has occurred in the past 42 Mya. Similar to exon-2, exon-3 is entirely missing in three species and has a CR1 element insertion in fourteen species considered (see **Fig. 7** and **Supplementary_information/Repeatmasker_results**). However, exon-3 is intact in the helmeted guineafowl (*Numida meleagris*). Hence, the exon-3 CR1 element insertion occurred sometime between 45 and 46 Mya, long after the gene loss event. In addition to the CR1 insertion in the exons, we found several CR1 insertion events within the *C9* locus of Galliformes. A comparison of CR1 presence in the *C9* locus across galliform species identified the presence of a CR1 insertion event at the approximate position of the erstwhile exon-10 shared by 16 of the 18 galliform species considered. This CR1 insertion at exon-10 is estimated to have occurred sometime between 45 and 46 Mya. Interestingly, we found a second CR1 insertion overlapping this exon-10-proximate CR1 in 12 of the 18 species considered. This second CR1 insertion spanned the exon-11 region and was potentially inserted sometime between 42 and 45 Mya. Compared to Palaeognathae, the prevalence of CR1 elements has increased at the *C9* locus in Neognathae (see **Supplementary_information/Repeatmasker_results**). However, the *C9* gene is intact in all non-galliform bird genomes considered and is transcriptionally active in representative species of Palaeognathae and non-galliform Neognathae.

## Discussion

The TCP has a prominent role in pathogen cytolysis. These TCP components are the result of gene duplication events. Gene synteny and phylogenetic analysis support the sequential duplication of the early C6-like genes through whole-genome duplication events (Mondragón-Palomino et al., 1999). The MACPF domain-containing C6-like proteins seen in cephalochordates and urochordates result from the divergence of perforin and early C6-like genes following the duplication of the MPEG-1 gene (D’angelo et al., 2012). The vertebrate *C6* contains CCP, FIMAC/KAZAL domains not found in any early *C6*-like proteins. Neither ancestral forms which contain all these protein domains nor other intermediate forms have been identified in genomic datasets. Hence, the birth of the vertebrate TCP components from earlier forms is not well understood. In contrast to the secondary loss evolutionary model (Peng et al., 2022) that explains the evolution of *C3, C4*, and *C5* genes, the birth of the vertebrate *C6* gene is hard to explain without invoking protein domain gain/gene fusion events. We found limited support when we considered a scenario of secondary domain loss (scenario-1 in Fig. 1A) for the emergence of vertebrate TCP. Similarly, scenario-3 (see **Fig. 1C**) involving independent domain gain by the different TCP components found limited support, and a post-WGD expansion of the vertebrate TCP is favored.

Two scenarios (see **Fig. 1B** and **Fig. 1D**) are consistent with this post-WGD expansion of the vertebrate TCP. While the second scenario (**Fig. 1B**) suggests a role for gene fusion, the fourth scenario (**Fig. 1D**) supports multiple sequential domain gain events. Comparative genomic approaches have helped us understand the mechanisms of protein domain gain during evolution (Buljan et al., 2010). Although retroposition, gene fusion, non-allelic homologous recombination, and illegitimate recombination have been proposed as mechanisms for domain gain, conclusively establishing the mechanism involved in any particular domain gain event is challenging. Hence, gene fusion is one of the major mechanisms for protein domain gain and is easier to establish based on shared exon phase, length, domain composition, and order. The gain of individual domains through gene fusion may be harder to identify due to the shorter size of the region under consideration. Therefore, our proposed second (**Fig. 1B**) and fourth (**Fig. 1D**) scenarios may not be mutually exclusive. Instead, they may represent a similar process of domain gain.

Certain domain combinations evolve more often than others and may result from parallel evolution (Zmasek and Godzik, 2012). The domain combinations found in TCP components are relatively novel and don’t occur in other proteins. However, the individual domains and combinations of two domains at a time occur in several proteins. For instance, the TSP1-LDLRA domain combinations occur in several mollusk genes. Similarly, the MACPF-EGF domain combinations are present in *PRF1*, and the CCP-FIMAC/KAZAL combinations occur in several genes. Hence, the multi-domain proteins of the TCP are potentially assembled gradually by sequential domain gain/gene fusion events. Protein domain gain is favored after donor/recipient gene duplication (Buljan et al., 2010). In the case of the *C6* gene, duplication of the recipient gene occurs before the gain of most protein domains (see **Fig. 3**).

**Figure 3:**
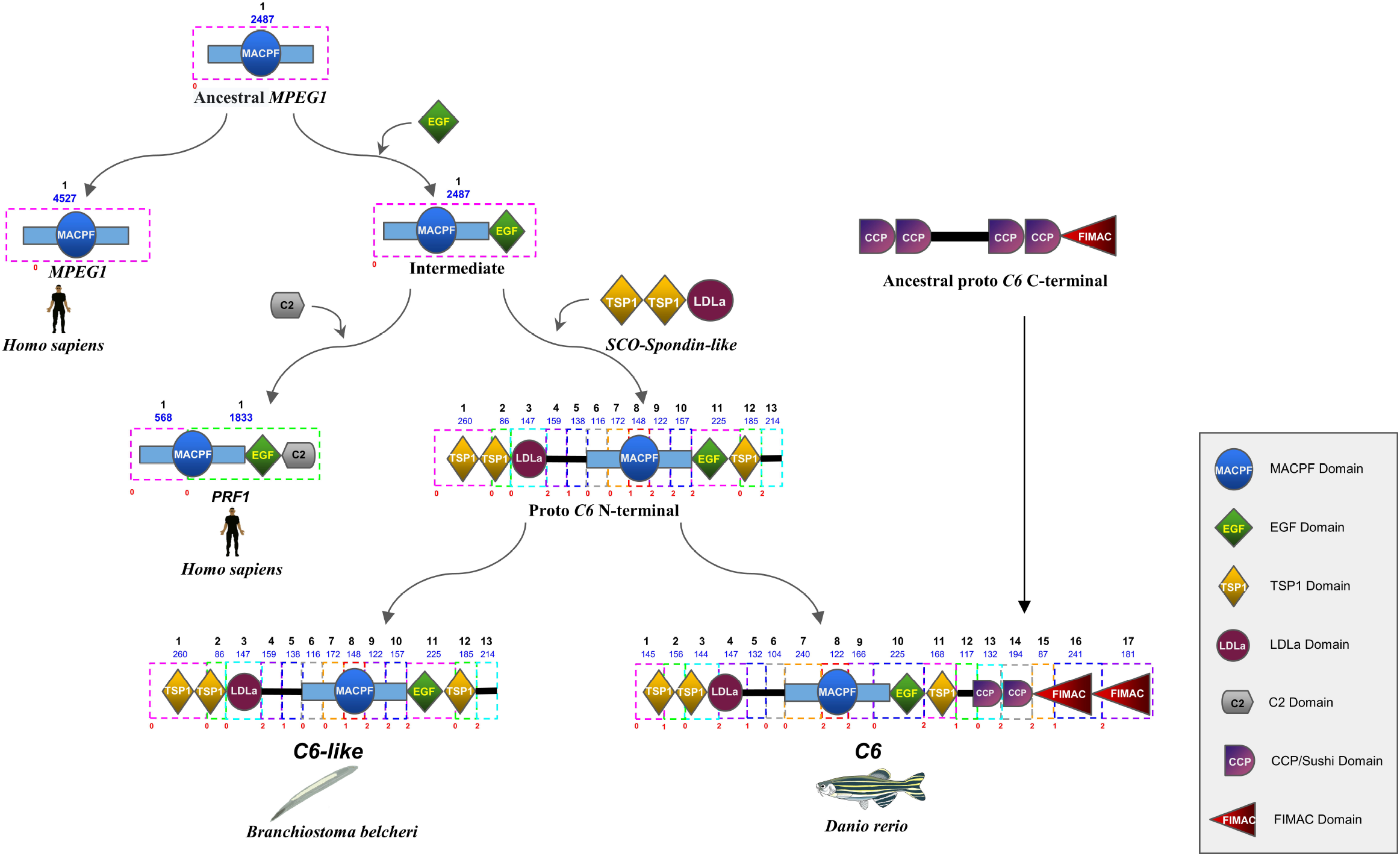
Schematic representation of the birth of late or terminal complement pathway (TCP) components: The emergence of vertebrate *C6* through a series of domain gain events is identified based on domain sequence similarity and matching exon length and phase. The dotted rectangular boxes around the different domains represent the exons. The exon rank and length are given above each box in black and blue, respectively. Each exon’s start and end phase are shown below the exon boxes in red color. Directional arrows represent the addition of domains. The solid black lines within genes represent the exonic sequences that did not match any protein domains.

**Figure 4:**
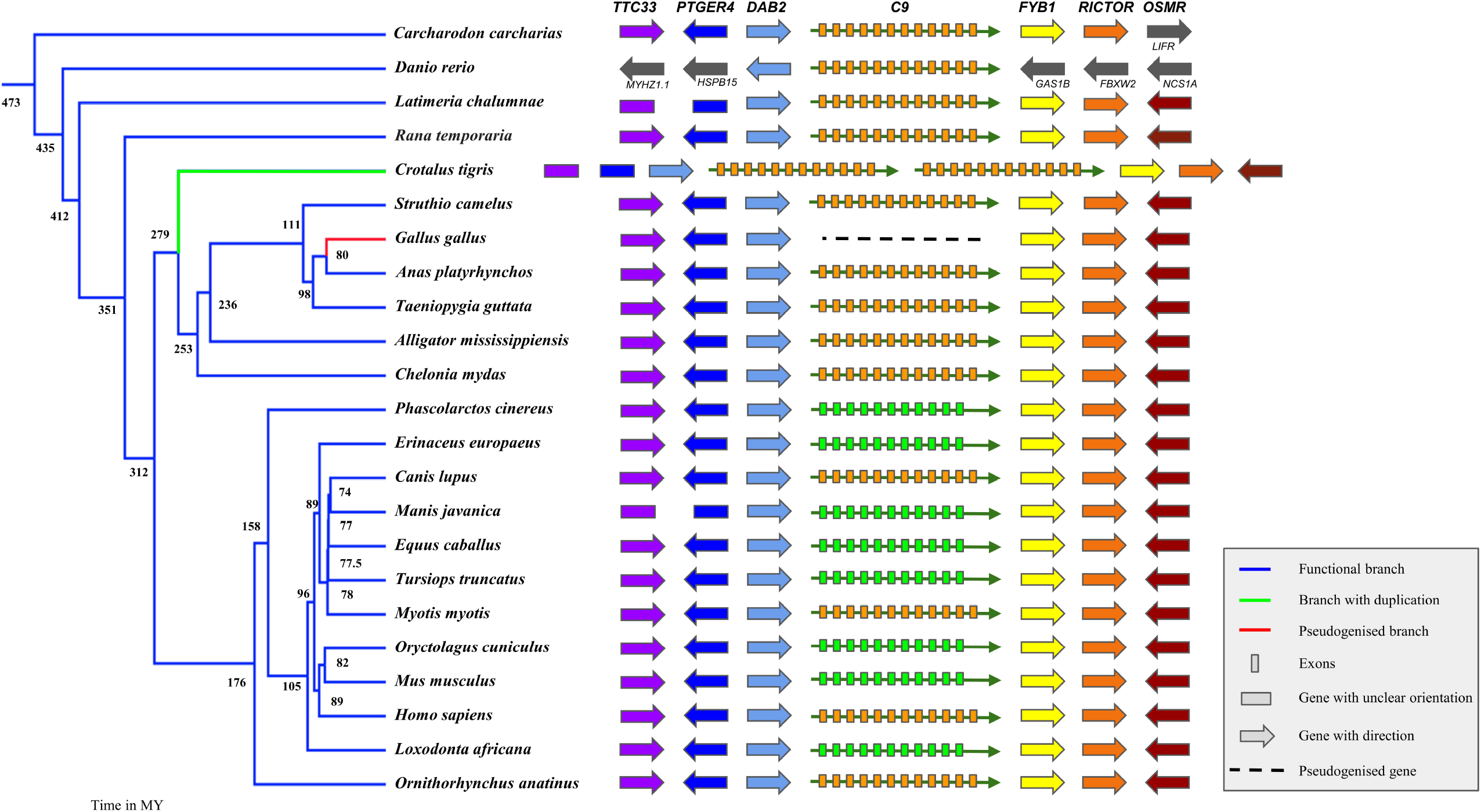
*C9* gene synteny in major vertebrate clades. Three genes upstream and downstream flanking the *C9* gene are shown. The arrowheads represent the gene orientation. Flanking genes represented by filled rectangular boxes are located on another scaffold, indicating unclear gene orientation. Genes found only in zebrafish (*Danio rerio*) and the great white shark (*Carcharodon carcharias*) are shown in dark grey color. A solid dark green arrow represents the intact *C9* gene, and a black dashed line represents *C9* gene remnants. Neighboring genes are colored in different colors, with each ortholog indicated in the same color. The conserved upstream genes are *DAB2* (Disabled homolog 2), PTGER*4* (Prostaglandin E Receptor 4), and *TTC33* (Tetratricopeptide Repeat Domain 33), while downstream genes are *FYB1* (FYN binding protein 1), *RICTOR* (Rapamycin-insensitive companion of mammalian target of rapamycin) and *OSMR* (Oncostatin M Receptor). *C9* gene duplication is shown by a green-colored branch in the phylogenetic tree, while the red-colored branch indicates a loss. Colored boxes indicate the exon structure of *C9*, with the number of boxes representing the number of exons in *C9. C9* copies containing eleven exons are shown with orange-colored boxes, while light green boxes represent ten exon copies.

**Figure 5:**
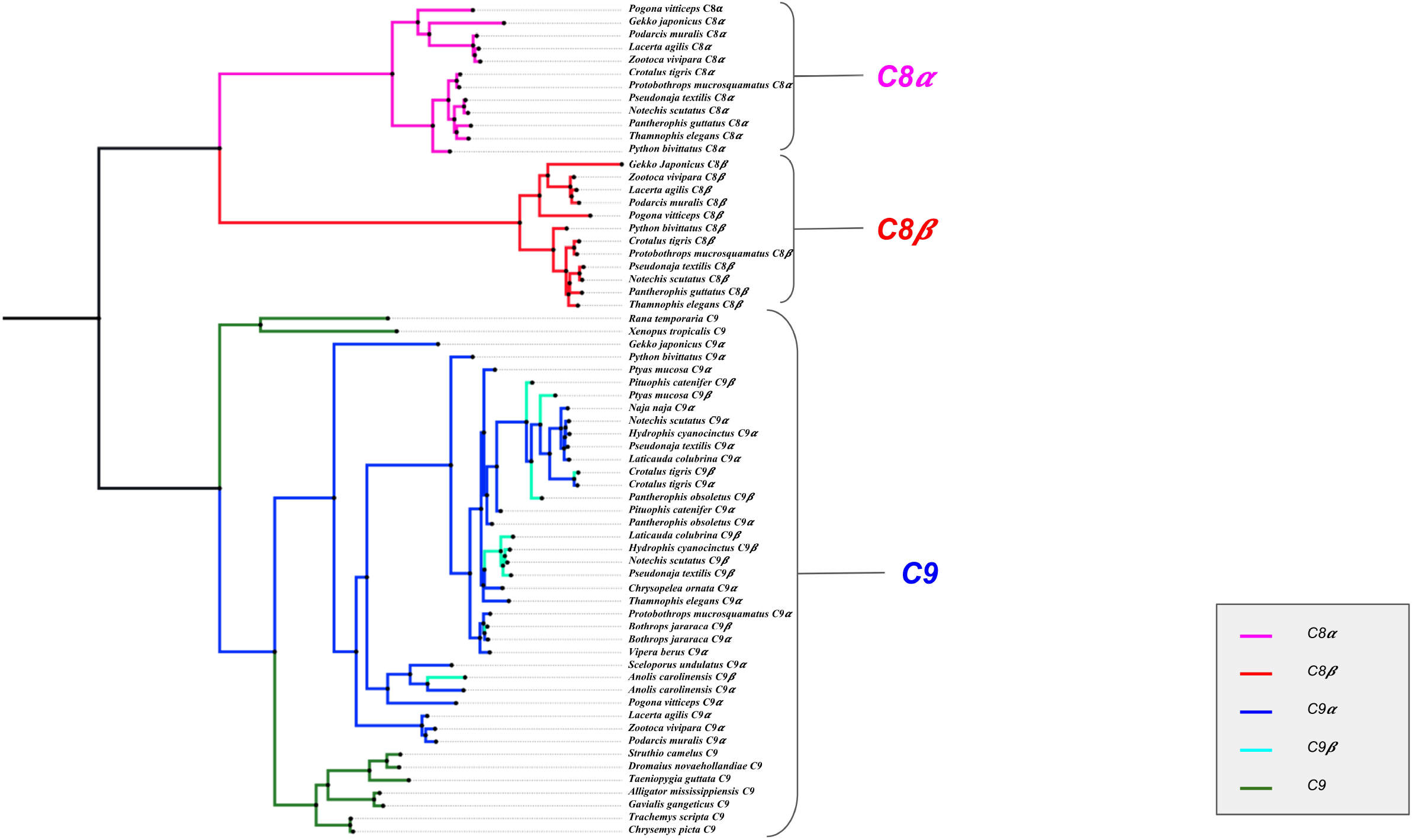
Phylogenetic relationship of *C8****α***, *C8****β***, and *C9* genes in Squamata. The maximum likelihood tree is generated from manually curated full-length open reading frames of *C8****α***, *C8****β***, and *C9* orthologs along with the two *C9* copies (i.e., *C9****α*** and *C9****β***) present in a few squamate species. The *C9* gene copy closer to the *DAB2* gene along the chromosome is referred to as *C9****α***, and the other copy closer to *FYB1 is* referred to as *C9****β***. Multiple sequence alignments used for building the tree are provided in **Supplementary_information/Fig5**. Pink branches and red branches represent *C8****α*** and *C8****β*** genes, respectively. The two copies of *C9* are shown in blue (*C9****α***) and cyan (*C9****β***) colors. Species with a single copy of the *C9* are shown in dark green color (for example, *Rana temporaria* and *Xenopus tropicalis*) and serve as outgroup sequences for the copies *C8****α*** and *C8****β***. Bootstrap support values are shown for each node in the **Supplementary_information/Fig5**.

**Figure 6:**
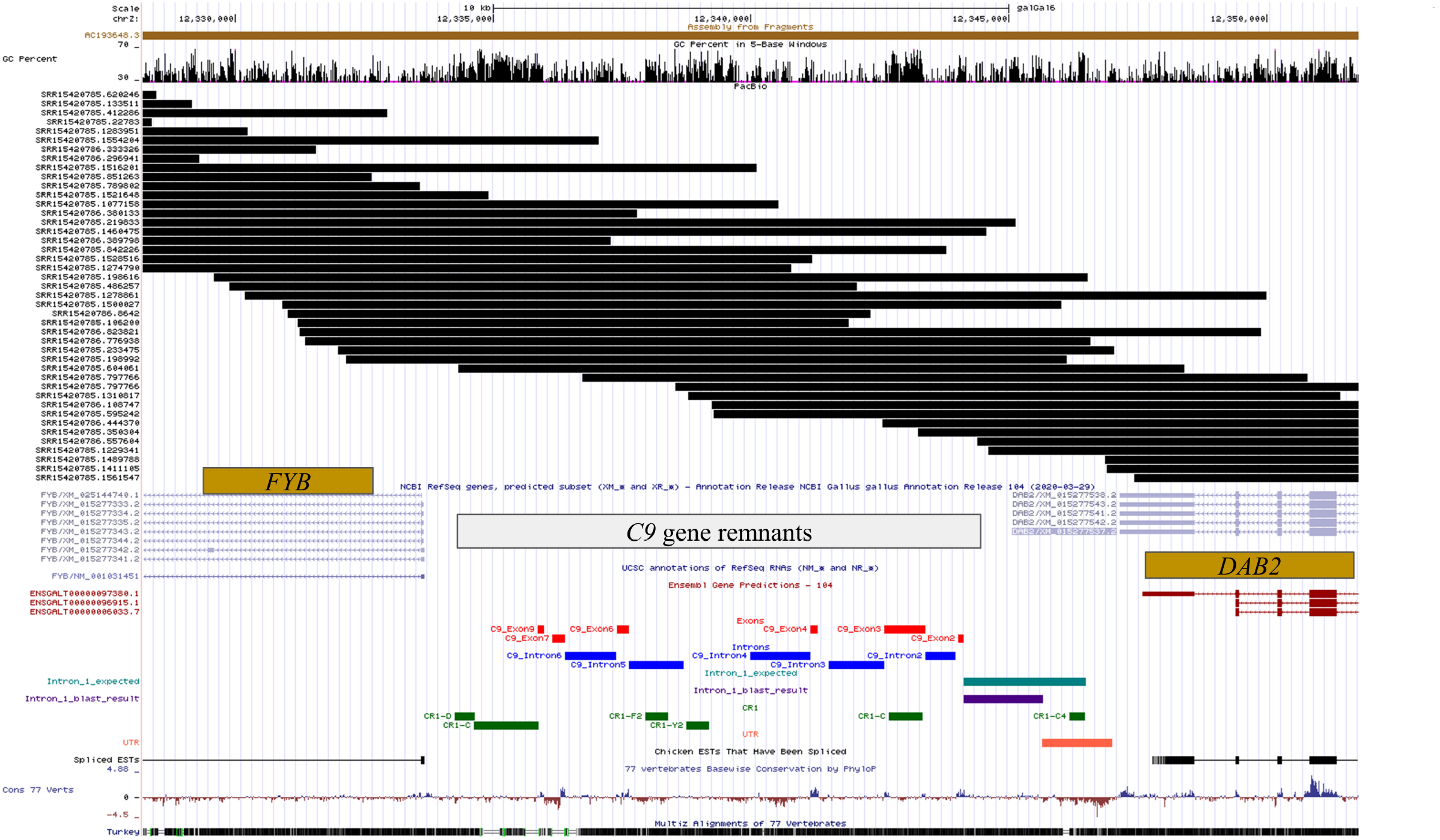
Verification of chicken genome assembly using PacBio long reads. Screenshot of UCSC genome browser showing the alignment of >10 kb long reads generated using PacBio technology and aligned to the chicken genome (GRCg6a) at the location of the *C9* gene remains. Red color boxes depict remnants of *C9* exons, and blue boxes represent the *C9* introns identified based on the *C9* gene sequence of the duck. The UTR, intron-1, and expected intron-1 regions are determined based on the duck *C9* gene sequence and shown by orange, purple, and dark-cyan colored boxes, respectively. Dark-green boxes represent the locations of the CR1 elements. Genes annotated in the chicken genome are represented by dark-yellow boxes (depicting exons) connected by thin lines (depicting introns). Each black rectangular box represents a read that aligns to that particular position. The accession numbers of each of the reads are provided beside the alignments. A series of overlapping reads span the entire region up to adjoining genes (*FYB* and *DAB2*) to form a tiling path.

**Figure 7:**
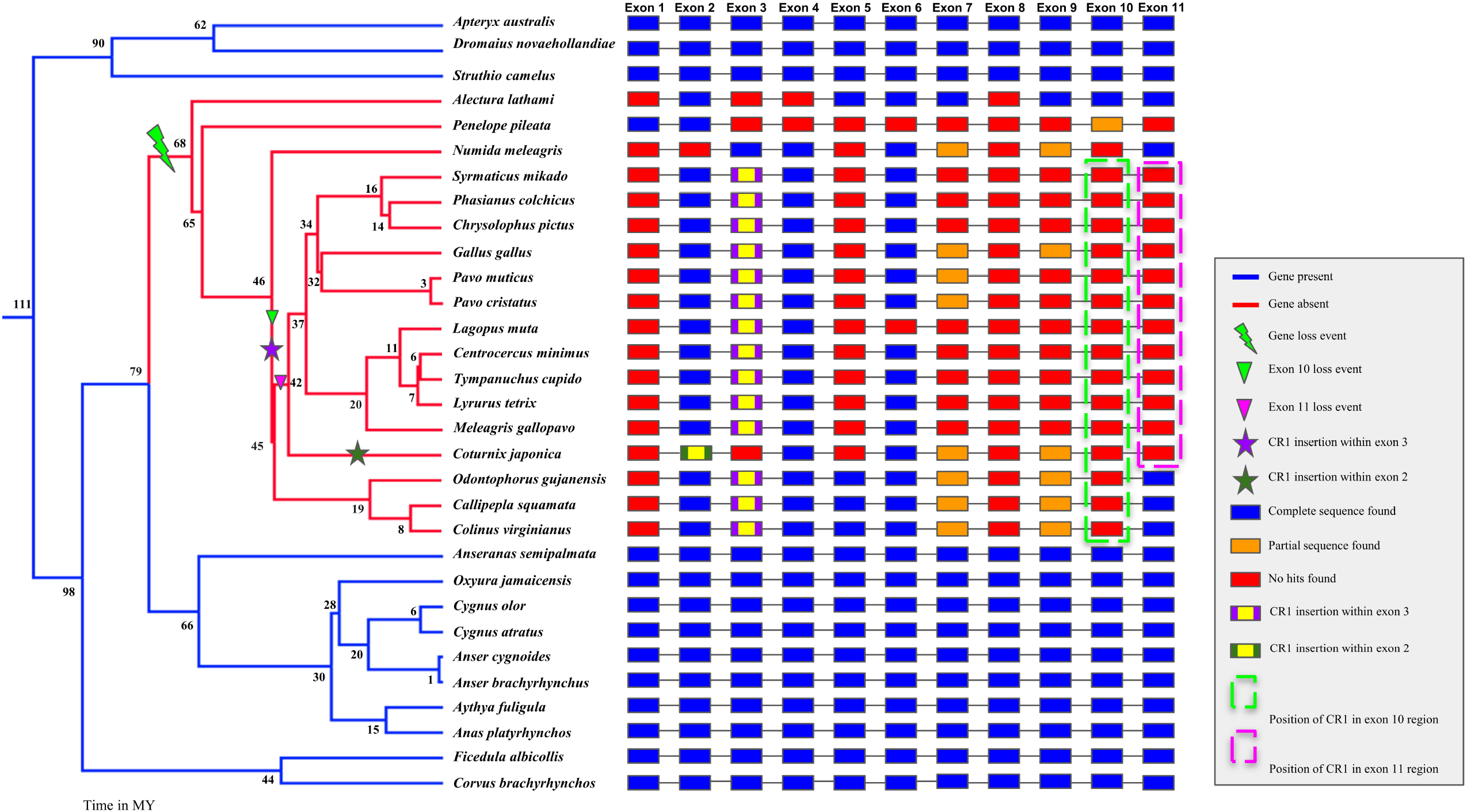
Loss of Complement component 9 (*C9*) in Galliformes. Loss of *C9* in 18 galliform species is shown along with their time-calibrated phylogenetic tree. Blue branches indicate the species with functional *C9*, while red branches indicate the species that have lost the *C9*. The green thunderbolt symbol shows a shared loss of exon eight in all 18 galliform species and potentially reflects the *C9* gene loss event. The purple “star” shows a CR1 (chicken repeat 1) insertion event in the third exon, and the green “star” shows CR1 insertion in the second exon of *Coturnix japonica*. The filled green and magenta triangles represent the loss of exon ten and exon eleven, respectively. Filled rectangular boxes represent the exons, and the lines connecting these boxes represent the introns. Blue color boxes represent the intact exon sequences, orange color boxes represent the exons with partial sequences, and red boxes represent the exon with no remnants. The purple rectangle with a yellow color box within represents a CR1 insertion in exon three. The green rectangle with a yellow-colored box within represents a CR1 insertion in exon two in *Coturnix japonica* only. Dotted rectangular green and magenta boxes represent the position of CR1 present in the region of exon ten and exon eleven, respectively.

The decreasing sequencing costs have led to ambitious projects that aim to generate genome assemblies of all living species (Lewin et al., 2018). The availability of such a large number of genomes spanning diverse evolutionary timescales will allow further scrutiny of the events that resulted in the birth of the TCP. Our study is limited by the number of high-quality genomes currently available. Future studies that have access to larger numbers of genomes would allow a higher resolution study of the sequence of events predicted to have given birth to the early components of the TCP. The proto-TCP proteins from Cephalochordata and Urochordata are an essential link in the birth of the modern TCP. Hence, experimental studies investigating the re-wiring of the complement pathway would help better understand the function of the proto-TCP. However, such functional studies would need to be conducted in non-model species and are beyond our study’s scope. Although our results cannot explain why the vertebrate TCP was favored, our analysis supports a sequence of events explaining how protein domain gains formed the proto-TCP.

A better understanding of immune system genes lost in specific evolutionary lineages and how it affects the host’s interaction with pathogens may provide important insights into combating diverse pathogens. Hence, the availability of high-quality genomes for many species has stimulated an interest in identifying lineage-specific gene loss events (Sharma et al., 2018; V. Sharma et al., 2020). The recent availability of a large number of bird genomes (Zhang et al., 2014a) and improved quality of the representative genomes from several clades of birds (Warren et al., 2017; Zhu et al., 2021) have allowed the detailed comparative investigation of bird genomes (Ellegren, 2013). For instance, the loss of *RIG-I* in the chicken while being functional in the duck has provided insights into the differential response of these bird species to various pathogens (Barber et al., 2010). The increasing availability of avian genomes has permitted the identification of recurrent loss of the *MDA5* gene, another cytosolic receptor involved in sensing viral RNA (Krchlíková et al., 2021). Several other genes involved in the immune response are potentially missing in birds (Magor et al., 2013). Identifying such natural gene knockouts has great potential in improving our understanding of basic biology and medicine (Emerling et al., 2017). Loss of several widely conserved genes in the Galliformes lineage while being intact in the closely related Anseriformes lineage has been identified (Sharad Shinde et al., 2021; S. Sharma et al., 2020). Other lineage-specific gene loss events within specific clades of birds have also been identified (Fiddaman et al., 2022; Friocourt et al., 2017; Haimson et al., 2021; Huang et al., 2022; Wicher and Fries, 2006).

Chicken belongs to the order Galliformes and has been a model organism for understanding immune gene functions and has contributed to several important discoveries (Kaufman, 2018, 2013; Parker and Kaufman, 2017; Tregaskes and Kaufman, 2021). Galliformes (land fowl) are primarily ground-dwelling, and the Anseriformes (waterfowl) live at the water surface. Changes in the pathogen repertoire between these groups of birds may result from the transition between land and water. This change in pathogen repertoire could explain the changes in the immune system genes and their regulatory mechanisms. Such pathogen load and content changes have occurred with the evolution of flight and blood-feeding in bat species (Banerjee et al., 2020; Blumer et al., 2022). Bat species have adapted to this pathogen load change by losing specific immune genes (Blumer et al., 2022). In the case of galliform birds, reports confirming the presence of Neisseria in the chicken are missing despite a large number of studies (Liu et al., 2015). Hence, the loss of the *C9* gene in galliform birds may be a consequence of relaxed selection resulting from a lack of infection from Neisseria pathogens. The lack of Neisseria infection in chicken will need to be investigated and may involve other host response mechanisms against Neisseria. Complement-resistant E. *coli* isolates express Lipopolysaccharide (LPS) O-antigen (O-Ag) to impair the polymerization of *C9* (Doorduijn et al., 2021). Recurrent infection by such complement-resistant strains could also be responsible for the relaxed selective constraints on the *C9* gene in galliform birds. In light of such recurrent infections, loss of the *C9* gene might provide a selective advantage against the harmful consequences of septic shock (Fu et al., 2016).

Galliform birds have lost genes from the plasminogen pathway (Beauclair et al., 2019; Chana-Muñoz et al., 2019; Daković et al., 2014; Leth and Ploug, 2021; S. Sharma et al., 2020). One of these genes, *PAI-1* (Plasminogen Activator Inhibitor 1), binds vitronectin (*VTN*) to modulate fibrinolysis and cell migration (Zhou et al., 2003). Hence, the loss of *PAI-1*, which serves as a switch controlling the interactions of *VTN*, would either lead to constant interaction or lack of interaction. *VTN*, along with clusterin (*CLU*), prevents the *C9* polymerization of the MAC (McDonald and Nelsestuen, 1997). Since the *PAI-1* gene is lost in galliform birds, the interaction of *VTN* with the MAC may be impaired and either entirely prevents or allows unregulated *C9* polymerization of the MAC. Hence, the loss of the *C9* gene in galliform birds might be a consequence of compromised regulation of *VTN*. The crosstalk between complement and coagulation pathways has recently witnessed considerable interest (Kenawy et al., 2015). Functional studies investigating the process of MAC formation in galliform birds might help better understand the role of *PAI-1* loss, *VTN* regulation, and polymerization of the MAC.

Paralogs or other members of the same pathway can take over the functions of a lost gene. Perforin is a distant homolog of the TCP components and has strong structural homology with *C9* at conserved regions (Shinkai et al., 1988). The pores formed by perforin are similar to those formed by the MAC and perform cytolytic activities (Baran et al., 2009). However, while the MAC acts against gram-negative bacteria and specific pathogenic parasites, perforin targets transformed host cells and cells infected by viruses (Kondos et al., 2010). Other components of the MAC may have adapted to the loss of *C9* to increase the efficiency of the *C9* deficient MAC. For instance, several lineage-specific amino acid changes in complement components *C5* and *C8****α*** have been reported in the Indian peacock (Jaiswal et al., 2018). How these lineage-specific changes affect function will require further studies. Functional characterization of the *C9* deficient MAC in galliform species would be the first step to understanding its abilities. In addition to the well-established function of cytolysis, other roles of TCP components are still being uncovered (Tegla et al., 2011). For example, TCP components *C9* and *C7* have been implicated as biomarkers for longevity in older men (Orwoll et al., 2020). Post-translation modifications of the complement *C9* protein result in several proteoforms which may have distinct functions (Franc et al., 2017). Being one of the most abundant proteins in the drusen (Crabb et al., 2002; Hollborn et al., 2018), complement *C9* is also referred to as *ARMD15* (Age-Related Macular Degeneration 15) and has long been associated with Age-Related macular degeneration (Kremlitzka et al., 2018; Zipfel et al., 2006) and multiple sclerosis (Morgan et al., 1984). The MAC has a complex interaction with cancer cells at different stages (Fishelson and Kirschfink, 2019), and elevated levels of *C9* have also been associated with esophageal adenocarcinoma (Joshi et al., 2017; Kolka et al., 2022) and colorectal cancer (Chantaraamporn et al., 2020). The complement system has considerable crosstalk with integrins to regulate single-cell metabolism (Merle et al., 2021). It will be intriguing to understand how the loss of *C9* in galliform birds affects these other functions.

The co-occurrence of CR1 insertions and gene loss events could be due to CR1 insertion mediated gene loss. However, the co-occurrence may simply reflect the mutational properties of a genomic region (Huang et al., 2022). Lineage-specific CR1 insertion also occurs in genes under relaxed purifying selection which have undergone recurrent gene loss (S. Sharma et al., 2020). Reconstructing the chronology of events that lead to gene loss will help understand the reason for the co-occurrence of CR1 insertion and gene loss events. In the case of galliform-specific *C9* gene loss, we found several CR1 element insertions. While most of these CR1 elements have accumulated after the *C9* gene loss, the reason for the complete absence of exon-8 in all galliform species is not clear. Non-allelic homologous recombination between CR1 copies can delete sequences resulting in the loss of exons (González and Petrov, 2012). Given the prevalence of CR1 elements at the *C9* locus, exon-8 deletion may result from recombination between CR1 elements flanking it. CR1 elements accumulate on the Z sex chromosome in birds (Bertocchi et al., 2018; Xu et al., 2019), and the presence of the *C9* locus on the Z chromosome in birds may explain the co-occurrence of CR1 elements with gene loss. Identifying all lineage-specific gene loss events and reconstructing the sequence of gene disruptive changes using high-quality complete genomes will allow for a comprehensive evaluation of the role of CR1 elements in gene loss.

## Conclusions

Based on the protein domain content of *C6*-like genes in pre-vertebrate species and comparing them to the protein domains found in vertebrate *C6* orthologs, we propose a hypothetical chronology of events that gave birth to the vertebrate *C6* gene. Our model suggests that the vertebrate *C6* gene results from sequential protein domain gain/gene fusion events. We used currently available genomic datasets to identify the genes that are potential donors of each domain based on sequence similarity, exon length, and phase. We find that the MACPF and EGF domains are obtained from an ancestral *MPEG-1* gene duplication event. A subsequent gene fusion between this intermediate and an SCO-Spondin-like gene resulted in an N-terminal *C6*-like gene. Finally, the vertebrate *C6* gene emerged from a second gene fusion event between N-terminal and C-terminal *C6*-like genes.

We use multiple independent genomic technologies to prove that the complement *C9* gene is disrupted in galliform birds. Detailed comparative genomic analysis of the *C9* locus identified several CR1 elements, including two within the exons. However, the other components of the TCP (*C5, C6, C7, C8****α***, *C8****β***, and *C8****γ***) are intact and transcriptionally active in galliform birds. Further studies will need to investigate how this galliform C5b-8 complex is different from other vertebrate species and why this *C9* gene loss event has occurred. Our detailed analysis of TCP genes in more than 350 species helps track the birth of the TCP components in the LVCA (Last Vertebrate Common Ancestor), their expansion through gene duplication events, and the death of the complement *C9* gene in galliform species. A similar study of other genes to unlock the mysteries of birth and death can provide helpful insight into gene function and adaptation.

## Acknowledgment

We thank the University Grants Commission for supporting AS with a Ph.D. scholarship and the Ministry of Human Resource Development fellowship to SG and ABP. The Department of Biotechnology, Ministry of Science and Technology, India (Grant no. BT/11/IYBA/2018/03) and Science and Engineering Research Board (Grant no. ECR/2017/001430) provided funds used to generate primary sequencing data published in this article and computational resources (i.e., Har Gobind Khorana Computational Biology cluster) used.

## Author contributions

A.S. and N.V. wrote the manuscript with major inputs from S.G, and A.B.P. A.S analyzed the data along with S.G. and A.B.P. Specific analysis benefited from insightful inputs provided by A.B.P, who also generated all the new sequencing data presented in this study. All authors reviewed the manuscript.

## Availability of data

This study’s primary sequencing data are available from the SRA under BioProject# PRJEB51011. All **Supplementary information**, including other associated data and scripts used for analysis, are provided in an easy-to-browse format: **https://github.com/Ashu2195/C9**. Once the manuscript is accepted, the Supplementary information will be published as a Mendeley Dataset.

